# Global chemical modifications comparison of human plasma proteomes from two different age groups

**DOI:** 10.1101/2020.03.07.978049

**Authors:** Yongtao Liu, Mindi Zhao, Xuanzhen Pan, Youhe Gao

## Abstract

The chemical modification of proteins refers to the covalent group reaction involved in their amino acid residues or chain ends which, in turn, change the molecular structure and function of the proteins. There are many types of molecular modifications in the human plasma proteome, such as phosphorylation, methylation, and acetylation. In this study, two groups of human plasma proteome at different age groups (old and young) were used to perform a comparison of global chemical modifications, as determined by tandem mass spectrometry (MS/MS) combined with non-limiting modification identification algorithms. The sulfhydryl in the cysteine A total of 4 molecular modifications were found to have significant differences: the succinylation and phosphorylation modification of cysteine (Cys, C) and the modification of lysine (Lys, K) with threonine (Thr, T) were significantly higher in the old group than in the young group, while the carbamylation of lysine was lower in the young group. Cysteine residue is an important group for forming disulphide bonds and maintaining the structure of the protein. Differential cysteine-related sulfydryl modifications may cause structural and functional changes. Lysine is a basic amino acid, and the modification of its amino group will change the charge state of the protein, which may affect the structure and function of the protein. In summary, four types of protein chemical modifications and substitutes were found to be significantly different in the plasma proteome in different age groups and their probabilities of random generation are lower by passing random grouping test. We speculate that there is an increase in certain modified proteins in the blood of the old people which, in turn, changes the function of those proteins. This change may be one of the reasons why the old people are more likely than the young people to be at risk for age-related diseases, such as metabolic diseases, cerebral and cardiovascular diseases, and cancer.

## 1 INTRODUCTION

The biological processes of living things/organisms can perform and operate in mutual coordination and high-efficiency cooperation simultaneously, relying on the synthesis, catalysis and regulatory reactions in which proteins are involved. Proteins are biological macromolecule with complex structures, and the difference in the high-level structures of proteins determine their different biochemical activities, which can perform specific functions through a higher-level network formed by the combination of proteins (Aebersold & Mann, 2016). Chemical modification of proteins refers to the covalent group reaction of amino acid residues or chain ends. In general, a few changes in chemical structure do not affect the biological activity of proteins, which are called modification of nonessential parts of the protein. However, in most cases, changes in molecular structure will significantly change the physical and chemical properties of the protein, change the conformation of the protein, and make the activity of the protein vary, and then the function that it performs is changed (Sakamoto & Hamachi, 2019). Therefore, even if there is no change in the protein content level, but some small changes in the level of chemical modification, the function of the proteins will change significantly, which means that the chemical modification of the proteins enriches the different functions of the protein in another dimension. The effects of chemical modification of proteins on the functions of proteins are mainly shown in the following three aspects: 1. Even if a modification occurs, the functions of proteins will be affected. 2. The same type or different type of modifications of the different amino acids have different effects on the same protein’s function. 3. The same protein may undergo many types of chemical modification, which makes the biological process in which it participates a more variable and complex process (C. Wang et al., 2019). The main types of chemical modifications related to proteins are as follows: 1. Post-translational modifications (PTMs) refer to the chemical modification of a protein after translation (Minguez et al., 2012). The precursor of post-translationally modified protein often has no biological activity. Post-translational modification of a specific modified enzyme is usually required for it to become a functional mature proteins and perform its specific biological functions (Goto, Kudo, Lee, & Oe, 2015; Pieroni et al., 2020). 2. Chemical-derived modification of proteins refers to a kind of modification that introduces new groups or removes original/intrinsic groups in the protein side-chain, usually including spontaneous non-enzymatic modifications, as well as the modifications introduced by cross-linking agents and artificial agents. 3. Amino acid substitution refers to the change in protein properties and functions caused by the substitution of amino acids in the protein side-chain by other kinds of amino acids. These changes are the types of modifications that significantly affect protein function.

Mass spectrometry can not only realize the acquisition of large-scale data and in-depth mining but also achieve the accurate determination of specific protein targeted modifications. With the continuous development of scientific instruments, ultrahigh resolution and tandem mass spectrometry (MS/MS) provide more abundant information or data for proteomics and chemical modification research, which also facilitates the accurate identification of chemical modification sites in the protein chain. In the process of proteomic data analysis, it is usually necessary to search and compare the proteomic database of species, and high-resolution tandem mass spectrometry is the primary method for obtaining a large amount of protein modification information. When using the search engine to search the database, known types of protein chemical modification were usually set. This type of search method is called a restricted search, but it is difficult to identify a new type of modification with a type in the product that is unknown (An et al., 2019). Therefore, comprehensive and non-limiting modification identification plays an important role in understanding all the chemical modification information contained in the sample proteome. Open-pFind is an open sequence library search algorithm that integrates the UniMod database, analyses and processes the collected mass spectrum data through an open search to obtain the global chemical modification information of samples (Chi et al., 2015; D. Q. Li et al., 2005; L. H. Wang et al., 2007).

Plasma is an important part of the internal environment and homeostasis, and it plays a role in transporting the substances needed for maintaining the life activities and the wastes generated off the body. Proteins are rich in variety and content in the plasma, and the components are easily affected by metabolism, physiology, and pathology. All metabolites or wastes in cells and tissues are transported and exchanged through the blood; the study of plasma proteomics may reflect the physiological state of the body at a specific stage. Chemical modification levels are another important research area of plasma proteomics. A comprehensive comparison of the changes in plasma proteome chemical modification levels will provide multiple dimensions of information for the study of physiological changes in the body (Goto et al., 2015). At present, there are more than 1500 kinds of chemical modifications in the UniMod (Creasy & Cottrell, 2004), PSI-MOD (Montecchi-Palazzi et al., 2008) and RESID databases. There are many kinds of chemical modifications in the human plasma proteome, such as N-terminal acetylation, phosphorylation in the side chain, methylation, glycosylation, ubiquitination and disulphide bonds between two chains. The plasma proteome can reflect the nuances defining the differences between age and ageing (Lehallier et al., 2019). A study of the proteome changes in another body fluid, urine, demonstrated that this fluid can be analysed to elucidate body ageing (X. D. Li & Gao, 2016). Even the common physiological process of hunger can be analysed to characterize the urine proteome (Yuan & Gao, 2017). However, the comparison of the global proteome chemical modifications in two kinds of body fluids (plasma and urine) showed the differences in modifications between different types of samples (Liu, Zhao, Pan, & Gao, 2020). Based on the importance of chemical modifications in the human plasma proteome, this study attempted to compare the differences in the global chemical modification levels of the plasma proteome at two different age groups, as determined by high-resolution tandem mass spectrometry combined with non-limiting modification identification (Open-pFind).

## 2 RESULTS

### 2.1 Identification of total protein by using the bottom-up proteomic technology

In the label-free proteomic analysis, 20 (10/10) samples were analysed by LC-MS/MS. After retrieving data (.raw) based on pFind studio, the analysis results can be browsed and exported in pBuild studio. The 120-minute liquid chromatogram gradient was analysed, and 628-781 kinds of proteins (average 724) and 7526-11464 kinds of peptides (average 10238) were identified in the plasma samples without high-abundance protein removal.

### 2.2 Comparison of post-translational modifications of the plasma proteome between the young and old groups

A total of 1169 modifications (including low abundance modifications, that is, each modification is identified at least once) were identified in the plasma samples of the old group, and 1154 modifications (including low abundance modifications) were identified in the plasma samples of the young group. Eighty-eight percent of the modifications are of the same type, and the remaining 12% are related (Figure 1).

**FIGURE 1.**
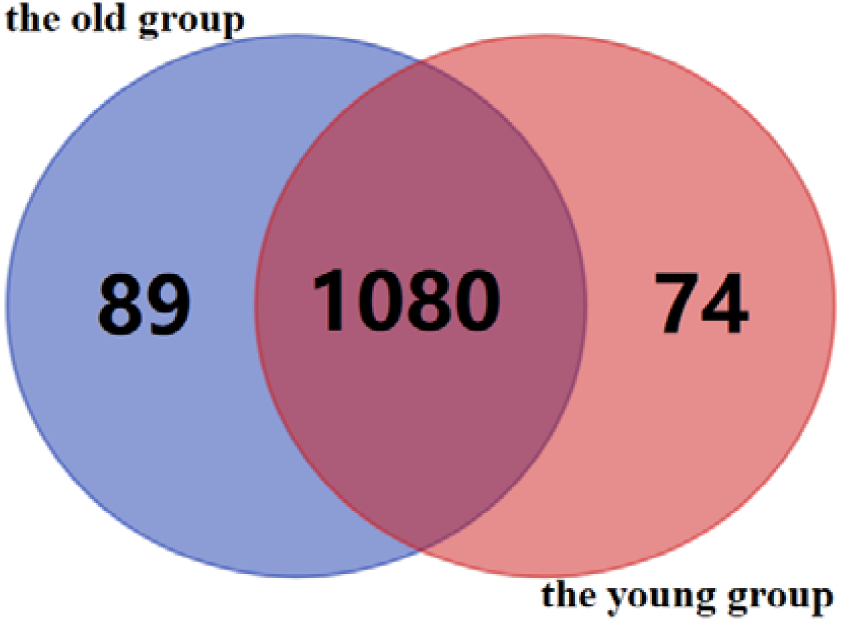
Statistics of the global plasma proteome chemical modifications in the old and young groups. The old and young groups have 1080 of the same type of chemical modification, of which 89 are unique to the old group, and 74 are unique to the young group. The above entire modifications include all low-abundance modifications.

Among the 163 unique protein chemical modifications in the two groups, 158 of them had less than 50% repeatability in each group, and only 5 of them met the conditions of more than 50% reproducibility. They are four kinds of modifications in the old group: Arg→Phe [R], Arg→Trp [R], Xlink_DMP [K], DTT_ST [S] and bisANS-sulphonate [T] modifications in the young group. In general, the unique chemical modification types are the molecular modifications with low identification quantity and poor reproducibility in the group.

Non-low-abundance chemical modifications (i.e., each modification is identified 10 times or more in the sample) are counted, and the identification coverage in each group is required to be greater than 50%. There are 120 kinds of chemical modifications in 1080 kinds that meet this condition. Unsupervised cluster analysis can roughly distinguish young group and old group samples, but 40% of young group and old group samples are clustered into one group, which is not classified as being the same as other samples in the group. Figure 2 shows the unsupervised clustering results of specific samples. The screening conditions for different chemical modifications are as follows: the p-value between two groups is less than 0.05, and the mean value of the number of chemical modifications of each sample in the group is calculated by the normalization of the total number of chemical modification spectra identified. There are four modifications with a multiple of change greater than 1.5 between groups: 2-succinyl [C], Lys→ Thr [k], phospho [C], and carbamyl [k]. Table 1 shows the details.

**TABLE 1.** Differential molecular modification information of plasma proteome between old and young group

| Order | No.* | Name | Type | p-value | Old avg | Young avg | Old/Young |
| --- | --- | --- | --- | --- | --- | --- | --- |
| 1 | 957 | 2-Succinyl[C] | Chemical<br>derivative<br>modification | 0.001 | 0.00091 | 0.00046 | 1.98↑ |
| 2 | 594 | Lys→Thr[K] | Amino acid<br>substitution | 0.003 | 0.00107 | 0.00065 | 1.65↑ |
| 3 | 21 | Phospho[C] | PTM | 0.013 | 0.00087 | 0.000548 | 1.59↑ |
| 4 | 1175 | Pro→Trp[P] | Amino acid<br>substitution | 0.042 | 0.00091 | 0.00061 | 1.49↑ |
| 5 | 21 | Phospho[S] | PTM | 0.027 | 0.00105 | 0.00077 | 1.36↑ |
| 6 | 64 | Succinyl[AnyN-term] | PTM | 0.032 | 0.00414 | 0.00317 | 1.31↑ |
| 7 | 23 | Dehydrated[S] | PTM | 0.022 | 0.0041 | 0.00446 | 1.09↓ |

*Note:* The number is the database entry number, from http://www.unimod.org/.

**FIGURE 2.**
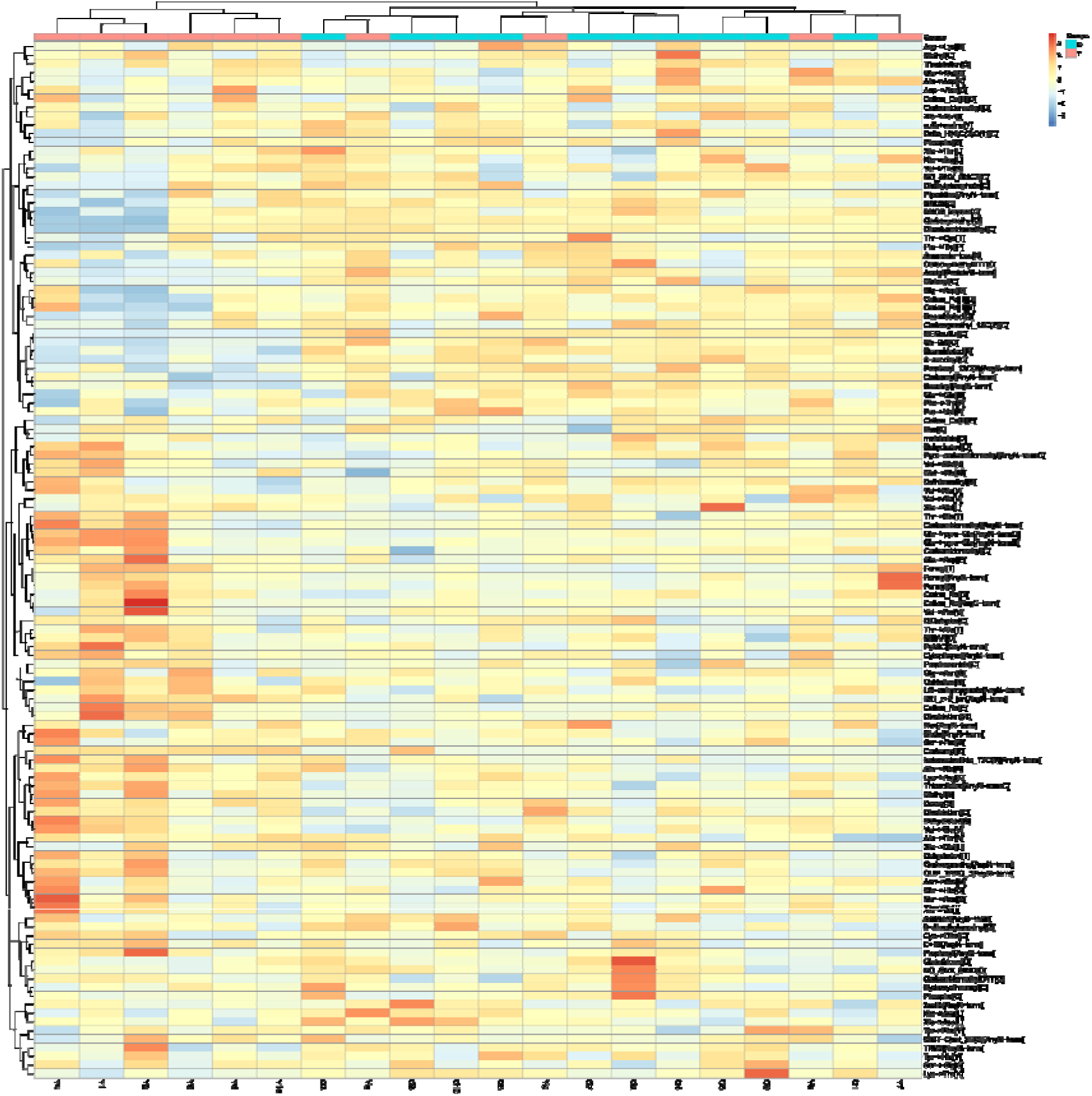
Unsupervised cluster analysis of overall plasma proteome molecular modifications in the young and old groups. The red mark (Y) is the sample of young group, and the blue mark (O) is the sample of old group. The abscissa is the unsupervised clustering and sample-specific information, and the ordinate is the specific molecular modification name.

### 2.3 Random grouping test

Four different modifications satisfying different screening conditions (p-value less than 0.05, fold change greater than 1.5) were performed by random grouping tests to verify the false-positive rate of each modification. Ten young group datasets and 10 old group datasets were randomly divided into two groups with 10 samples in each group and 92,378 different combinations 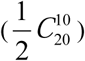. Each combination was statistically analysed by the same difference screening conditions. After extensive calculation and statistical analysis, four different modified random combinations were obtained. See Table 2 for details.

**TABLE 2.** Random grouping calculation results of 4 different chemical modifications in the plasma proteome

| No. | Name | Total times in random<br>groups | Frequency of<br>occurrence | Confidence level% |
| --- | --- | --- | --- | --- |
| 1 | 2-succinyl[C] | 4857 | 0.0525 | 94.74 |
| 2 | Lys→Thr[K] | 1135 | 0.0123 | 98.77 |
| 3 | Phospho[C] | 2126 | 0.0230 | 97.70 |
| 4 | Carbamyl[K] | 4664 | 0.0505 | 94.95 |

Through the random grouping test, it is found that the randomness of the four kinds of different modifications is approximately 5%, and the reliability is more than 90%. It is shown that the difference between the four kinds of modifications in different age groups is less likely to be generated at random.

## 3. CONCLUSIONS AND DISCUSSION

The entire life cycle of proteins, extending from translation assembly to final degradation, involves many toxic environments. These environments modify protein molecules to change the structure and function of the protein. This process is also called molecular ageing (Gillery & Jaisson, 2013). These reactions are mainly caused by the non-enzymatic binding of active small molecules on proteins with functional groups. The modification of proteins can be divided into reversible and irreversible modifications. Through a non-limiting modification search, several significantly different protein chemical modifications were found in the plasma of the two different age groups. These modifications include succinamide modification of cysteine, substitution of lysine residues by threonine residues, phosphorylation of cysteine, substitution of proline residues by tryptophan residues, phosphorylation of serine, N-terminal modification of succinamide, dehydration of serine, substitution of glutamic acid residues by aspartic acid residues, and carbamylation of lysine. Among these changes, succinylation, phosphorylation of cysteine and L-lysine replacement with threonine were significantly higher in the older group than in the younger group, and carbamylation of lysine was higher in the younger group than in the older group.

Cysteine is one of the amino acids with the lower abundance in the protein. The statistics of protein residues show that the average frequency of cysteine in eukaryotes is approximately 2%, and 70% of the reduced sulfhydryl source protein is present in the *in vivo* environment (Pe’er et al., 2004). Studies have shown that the abundance of cysteine in protein is affected by the function of the protein. The high activity of the sulfhydryl group makes facilitates its participation in many chemical modifications. In contrast, this group also determines the distribution and topological properties of cysteine residues in protein structure. The sulfhydryl group is also an important group to form a disulphide bond and maintain protein structure (Gunnoo & Madder, 2016).

The succinylation and phosphorylation of cysteine in this study involve the participation of the sulfhydryl group, which may affect the formation of the disulphide bond. Fumaric acid was added to the dissociative sulfhydryl sites of some Cys residues in proteins by a Michael addition reaction to form S-(2-succinic acid) cysteine (Alderson et al., 2006). This modification was initially detected in plasma proteins (including albumin) and formed by irreversible reactions. This modification has been reported in diabetes, obesity, fumaric acid hydratase-related diseases and the model of RIE’s syndrome. At the same time, it was also found that the content of succinic acid protein increased in mouse 3T3-L1 adipocytes cultured in high glucose medium (30 mm, while the physiological level was 5 mm), as well as in rats treated with streptozotocin (Blatnik, Thorpe, & Baynes, 2008; Frizzell et al., 2009; Thomas, Storey, Baynes, & Frizzell, 2012). It has been reported that an excess of nutrients (sugars) will lead to an increase in ATP: ADP, NADH: NAD^+^ and mitochondrial membrane potential, while an increase in NADH: NAD+ will inhibit oxidative phosphorylation, resulting in the continuous accumulation of mitochondrial intermediates (including fumaric acid), leading to an increase in protein succinylation (Zheng et al., 2015). The accumulation of succinate protein is also caused by a decrease in fumarate hydratase activity. Fumaric acid hydratase catalyses the reversal of fumaric acid to malic acid in the tricarboxylic acid cycle. It is important to note that loss of function and mutations in fumaric hydratase are known to predispose affected individuals to multiple skin diseases and uterine leiomyomas, as well as hereditary leiomyomas and renal cell carcinoma (HLRCC) (Pe’er et al., 2004). Although the exact role of succinic acid-modified cysteine and other related proteins has not been fully elucidated, it is related to the cancer response related to fumarate hydratase (Adam et al., 2011; Bardella et al., 2011; Ooi et al., 2011). The increase in protein succinic acid was also described in the brain stem of ndufs4 knockout mice (a model of Leigh syndrome) (Piroli et al., 2016), indicating that this type of protein chemical modification has a potential role in the pathogenesis of this mitochondrial disease. Park et al. found that in the detected protein succinylation sites, 16 succinylation sites appeared in the cofactor binding area or enzyme catalytic area, and 74 succinylation sites existed around the enzyme active site (Piroli et al., 2014). Baynes et al. found that cysteine succinyl modification in human skin collagen increased with age (Blatnik et al., 2008; Frizzell, Lima, & Baynes, 2011; Frizzell et al., 2009).

Cysteine phosphorylation was recently found in prokaryotic and eukaryotic systems and is believed to play a key role in signal transduction and regulation of cellular responses. Due to the low chemical stability of thiophosphates in peptides, the *in vivo* phosphorylation of Cys side chains is rarely studied(Attwood, Piggott, Zu, & Besant, 2007; Zu, Besant, Imhof, & Attwood, 2007). Phosphorylation of cysteine is an important function of cysteine-dependent protein phosphatase (CDP), which belongs to a subfamily of protein tyrosine phosphatases (PTPs) and catalyses the hydrolysis of phosphate ester bonds through the formation of phosphate cysteine intermediates. Studies have shown that this reversible post-translational regulation (PTM) is crucial in regulating the expression of virulence determinants and bacterial resistance to antibiotics. In addition, it has also been shown that phosphorylation of cysteine, previously considered a rare modification, may be more common in nature and may play an important role in the biological regulation of various organisms.

The replacement of lysine with threonine changed the acid-base properties of the protein, which affected the activity and function of the protein. It has long been observed that the substitution of lysine residues with threonine residues on haemoglobin will reduce its oxygen affinity (Barberio, Leone, Ivaldi, & Giordano, 2013). Sun WY et al. found that the mutation of nucleotide 20040 in exon 14, with Thr replacing Lys at amino acid 556, would reduce the pro-coagulatory activity of prothrombin by 50% (Sun, Smirnow, Jenkins, & Degen, 2001). The decrease in oxygen affinity, including some other recessions belonging to coagulation activity and other physiological conditions, also reflects the slow metabolism of the body, which may be the earlier manifestation of ageing. In addition, it was found that human and mouse embryonic stem cells need specific amino acids to proliferate. MES cells need threonine (Thr) metabolism to complete epigenetic histone modification. Thr is converted to glycine and acetyl coenzyme A, and glycine metabolism specifically regulates the trimethylation of lysine (Lys) residues in histone H3 (H3K4me3) (Van Winkle & Ryznar, 2019). In addition, we also found that the modification of L-lysine carbamylation in the old group was lower than that in the young group. Carbamylation is an irreversible non-enzyme modification process. The process is the side chain reaction between the decomposition product of urea and the N-terminus of protein or lysine residue, which was previously reported to be related to the ageing of proteins (Van Winkle & Ryznar, 2019). L-lysine carbamylation can promote the coordination interaction of metal ions to specific enzyme activities. Some studies have pointed out that the amount of carbamylation in the plasma of patients with increased urea levels (such as nephrotic patients) is significantly increased (Badar, Arif, & Alam, 2018).

Ageing is an inevitable and spontaneous process undergone by the organism with the passage of time. Ageing is a complex natural phenomenon that is manifested by the degeneration of structure, the decline of function, and the recession of adaptability and resistance. Ageing is one of the largest known risk factors for most human diseases: approximately two-thirds of the world’s 150000 people die every day from ageing-related causes. At present, the cause of ageing has not been determined. The current mainstream theory explaining ageing is damage theory, and DNA damage is considered to be the common foundation of cancer and ageing. Some people think that the internal cause of DNA damage is the most important driving force of ageing. According to the waste accumulation theory, the accumulation of waste in cells may interfere with metabolism. For example, a waste called lipofuscin is formed by a complex reaction of fat and protein combining in cells. These wastes accumulate in cells in the form of small particles, and their size will increase with age. Plasma protein can reflect changes in the body during the process of ageing (Lehallier et al., 2019). At the same time, the accumulation of biological macromolecules whose structure is destroyed or even inactivated may lead to the gradual failure of the biological body and system, which is also considered to be the concept of ageing.

Through the research described above, we found several types of chemical modifications and replacement of proteins in different age groups. These chemical modifications change the structure and properties of proteins and then affect the function of proteins. It is speculated that the gradual accumulation of some kinds of harmful and irreversible protein modifications in the plasma of the elderly may reflect the ageing process of the body. The accumulation of harmful protein modifications may be one of the reasons why the old are more likely than the young to suffer from ageing-related diseases, such as metabolic diseases, cardiovascular and cerebrovascular diseases and tumour risks.

## 4 EXPERIMENTAL PROCEDURES

### 4.1 Sample collection

The plasma samples of 20 patients were collected from the clinical laboratory of Beijing Hospital, and all of the samples had been discarded from the clinical laboratory. The samples were divided into two groups according to age: a young group and an old group. The samples were randomly selected from the clinical laboratory samples, and there were no restrictions or requirements on the diet, drugs and other factors of the blood sampling donors. The study was approved by the Beijing Hospital and the ethics committee of Beijing Normal University. This experiment provided volunteers with detailed information about the study, including its purpose, method, and process, and kept the personal information of volunteers strictly confidential. Only age and gender information are mentioned for discarded samples from the laboratory. See **TABLE 3** for sample information.

**TABLE 3.** Statistical data of plasma samples.

| Sample type | Plasma in the young group (n=10) | Urine in the old group (n=10) |
| --- | --- | --- |
| Gender: |  |  |
| Male | 2 | 3 |
| Female | 8 | 7 |
| Age: |  |  |
| Median | 31.8 ± 4.19 | 59.3 ± 3.46 |
| Average | 26-37 | 56-66 |

### 4.2 Protein sample preparation and trypsin enzymolysis

The plasma was centrifuged after whole blood anticoagulant treatment. The plasma sample (n = 20) was diluted 40 times with Milli-Q water, and then 100 μl was taken for subsequent experiments. A 20 mmol / L dithiothreitol (DTT) was used to react with the sample at 37°C for 1 h to denature the disulphide bond in the protein structure, and then 55 mmol / L iodoacetamide (IAM) was added and reacted in the dark for 30 min to alkylate the disulphide bond sites. The supernatant was precipitated with three volumes of precooled acetone at −20°C for 2 h and then centrifuged at 4°C and 12000 × g for 30 min to obtain protein precipitation. The precipitate was then resuspended in an appropriate amount of protein lysis buffer solution (8 mol / L urea, 2 mol / L thiourea, 25 mmol/L DTT and 50 mmol/L Tris). The concentration of protein extract was measured by Bradford analysis. Using filter-assisted sample preparation (FASP), trypsin gold (mass spec grade, Promega, Fitchburg, WI, USA) was used to hydrolyse 100 μg protein at a ratio of 50:1 for each sample. The dried peptides were sealed at −80°C after drying by a vacuum centrifugal concentrator.

### 4.3 Liquid chromatography tandem mass spectrometry (LC-MS / MS) analysis

Before analysis, the dried polypeptide samples were dissolved in 0.1% formic acid solution, and the final concentration was controlled at 0.1 μg/μL. Each sample was analysed according to 1 μg polypeptide quality: a Thermo Easy-nlc1200 chromatographic system was loaded on the precolumn and the analytical column. Proteomic data were collected by a Thermo Fisher Scientific (Bremen, Germany) mass spectrometry system. Liquid chromatography: pre-column: 75 μ m × 2 cm, nanoviper C18, 2 μ m, 100 Å; analytical column: 50 μ m × 15 cm, nanoviper C18, 2 μ m, 100 Å; injection volume: 10 μ L, flow rate: 250 nL/min, mobile phase: phase A: 100% mass spectrometry grade water (Fisher Scientific, span)/1‰ formic acid (Fisher Scientific), phase B: 80% acetonitrile (USA)/20% water/1‰ formic acid, 120 min gradient washing off: 0 min, 3% B phase; 0 min-3 min, 8% B phase; 3 min-93 min, 22% B phase; 93 min-113 min, 35% B phase; 113 min-120 min, B phase; mass spectrometry, ion source: spray voltage: 2.0kv, capillary temperature:320°C, resolution setting: first-order(Orbitrap) 120,000 @m/z 200, second-order (Orbitrap) 30,000 (Orbitrap) @m/z 200,parent ion scanning range: m/z 350-1350; sub ion scanning range: start from m/z 110, MS1 AGC: 4E5, charge range: 2-7, ion implantation time: 50 ms, MS 2 AGC: 1E5, ion implantation time: 50 ms, ion screening window: m/z 2.0, fragmentation mode: HCD, energy NCE 32, data dependent MS/MS: Top 20, dynamic exclusion time: 15 s, internal calibration mass number: 44 5.12003.

### 4.4 Database search

The pFind Studio software (version 3.1.3, Institute of Computing Technology, Chinese Academy of Sciences) was used to analyse the LC-MS / MS data with label-free quantification. The target retrieval database is from the Homo Sapiens database downloaded from UniProt (updated to October 2018). At the time of retrieval, the instrument type is HCD-FTMS, the full specificity of an enzyme is trypsin, and there are at most two missing sites. Open-search is selected. Screening conditions: FDR at the peptide level is less than 1%, and the Q value at the protein level is less than 1%. Data are analysed by using both forward and reverse database retrieval strategies.

## ACKNOWLEDGMENTS

We would like to thank Dr. Hao Chi for providing help in pFind Studio. This study was supported by the National Key Research and Development Program of China (2018YFC0910202, 2016YFC1306300), Beijing Natural Science Foundation (7172076), Beijing Cooperative Construction Project (110651103), Beijing Normal University (11100704), and Peking Union Medical College Hospital (2016-2.27).

## AUTHOR CONTRIBUTIONS

Y.L. designed and conducted the experiments, analyzed the data, prepared the figures, and wrote the paper. M.Z. provided samples and helped conducting the experiments. X.P. retrieved of references and helped with writing the paper. Y.G. provided the main ideas, designed experiments, and helped with writing the paper.

## DATA AVAILABILITY STATEMENT

The data that support the findings of this study are available from the corresponding author (Y.G), upon reasonable request.

